# Resolving FRET Signal Degeneracy and Population Heterogeneity via Bayesian Nonparametrics

**DOI:** 10.1101/2025.06.12.659382

**Authors:** Ayush Saurabh, Gde Bimananda Mahardika Wisna, Maxwell Schweiger, Rizal Hariadi, Steve Pressé

## Abstract

Biomolecular dynamics are often strikingly heterogeneous, with individual molecules sampling different states and kinetics—violating the “average molecule” assumption. Yet FRET analyses cannot resolve such variability or distinguish states differing mainly in kinetics, rather than FRET efficiency, as molecular configurations are projected onto 1D FRET signals. Here we introduce BNP-FRET-Bin, inferring state numbers and their kinetics directly from FRET data. In doing so, we eliminate user-specified parameters and expose molecule-to-molecule heterogeneity revealing new biologically relevant Holliday junction states with near identical FRET efficiencies.

## Main

Förster resonance energy transfer (FRET) is a nanometer-scale spectroscopic ruler that has illuminated intra- and intermolecular dynamics in proteins, nucleic acids, and their interactions [1]. Recent experiments have revealed that populations of single molecules probed by FRET can be highly heterogeneous [2, 3]: different molecules in the same sample access different numbers of conformational states, transition between those states at different rates, and form distinct subpopulations and rare pathways. Consequently, analyzing the data under the assumption that different traces sample an “average molecule” is misleading. The challenge is heightened by the fact that FRET collapses molecular configurations onto a 1D signal, making states that differ mainly in kinetics difficult to distinguish by FRET efficiency and analyses that rely on it. Both intrinsic noise, such as photon shot noise, and instrument-related contributions—including detector noise, background, spectral crosstalk, and nonideal fluorophore emission—further obscure state identity and kinetics. These combined effects place unusually high demands on analysis methods, yet widely used FRET tools often impose simplifying assumptions (for example, Gaussian noise, uncalibrated pixels, or fixed state counts) or require supervised training on simulated data—assumptions that are difficult to justify for heterogeneous, low-SNR datasets [1, 4, 5, 6].

These simplifying assumptions lead to concrete, predictable failure modes. First, inaccurate noise estimates can produce under- or over-fitting [7]. Second, because different traces visit different states and exhibit different kinetics, users must either choose the number of states separately for every trace—an impractical burden—or impose a single choice across all traces, which again enforces the notion of an “average molecule”. Third, incorrect determination of the number of global states across all traces creates internal inconsistencies; for example, identifying peaks in FRET-efficiency histograms can undercount states when distinct conformations yield similar or near identical efficiencies. Other model-selection criteria, such as BIC, AIC, and empirical Bayes (EB) [1], penalize model complexity but remain unreliable when data are sparse or when traces sample different subsets of states. In such settings, AIC/BIC may not sufficiently offset likelihood increases arising from adding states [8], and EB can return over- or under-confident priors depending on which subset of heterogeneous traces dominates. Moreover, because these approaches return only point estimates rather than full model uncertainty, they can be unstable to small data changes, prone to overfitting noise, and unable to capture multimodal solutions where different state combinations explain the data.

Bayesian nonparametrics (BNP) [9, 10] provides a unified framework for addressing inaccurate noise modeling, fixed-state assumptions, trace-to-trace differences in visited states, kinetic heterogeneity, and the lack of uncertainty quantification. Using BNPs, the number of states is treated as a random variable inferred jointly with kinetic rates and FRET efficiencies, enabling inference in an effectively infinite-dimensional state space. This flexibility suppresses unwarranted state proliferation when data are sparse because the prior disfavors unnecessary complexity—while avoiding underfitting by expressing appropriately large posterior uncertainty. The resulting posterior naturally reflects data quality, including signal-to-noise and trace length. BNP also resolves a central challenge of single-molecule datasets: different molecules may occupy different conformational states and transition among them at different rates, and imposing a shared state space (assuming an “average molecule”) across traces can mask genuine molecule-to-molecule differences. BNP instead allows each molecule to reveal the states it actually visits, preserving rare or molecule-specific behaviors that fixed-state or hierarchical approaches often suppress. Finally, BNP can identify states shared across some molecules, unique to others, or rarely visited, with all traces contributing jointly to the inference.

As such, we present BNP-FRET-Bin [11], a Julia package that provides a complete workflow consisting of: 1) a script to extract FRET traces from raw TIFF images recorded by sCMOS or EMCCD cameras; 2) a script to generate camera noise maps; and 3) an analysis tool implementing a nonparametric hidden Markov model (HMM) that produces Monte Carlo samples for all parameters of interest. For each trace, samples of the kinetic rate matrix, FRET efficiencies, and state trajectories are stored in a single HDF5 file. A companion script aggregates these files to generate probability and transition-density heatmaps.

To evaluate BNP-FRET-Bin, we first simulated a three-state FRET trajectory with realistic sCMOS noise characteristics (Poisson photon statistics followed by Gaussian readout noise), as shown in Fig. 1a–b. BNP-FRET-Bin correctly identifies the three-state model as most probable while assigning non-negligible probability to four- and five-state models (Fig. 1c–g), reflecting genuine uncertainty arising from intermediate signal-to-noise ratios and limited dwells in the highest and lowest FRET states. Our computational strategy (Methods) also yields sampled state trajectories that closely follow the apparent FRET signal. We then examined a scenario with FRET-efficiency degeneracy (Fig. 1h-i), where states 1–2 and 3–4 share similar efficiencies and differ only in kinetics. Histogram-based approaches infer only two states, which leads to rate averaging, misassignment, and overfitting. BNP-FRET-Bin instead explores models of varying dimensionality and recovers the kinetically distinct yet efficiency-degenerate states, along with their transition rates (Fig. 1j–n). We emphasize that discovering these states in the data is not possible from analyzing FRET efficiency alone.

**Figure 1:**
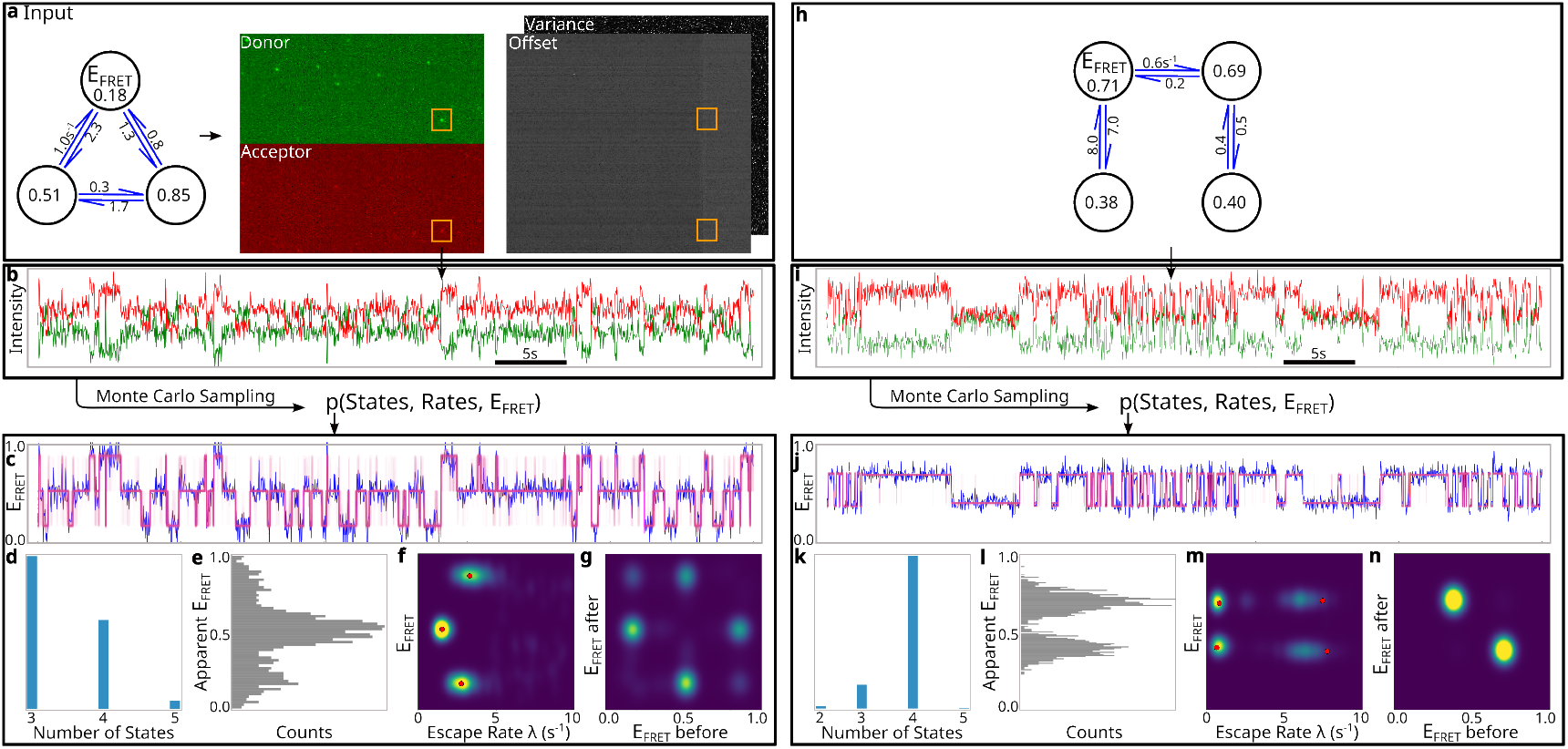
Schematic of the BNP-FRET-Bin algorithm. **a**. The kinetics of biomolecules are observed under a microscope using a camera. The two camera channels help visualize donor and acceptor fluorescence as diffraction-limited spots at the same locations (highlighted here using orange squares). Camera noise calibration maps for readout offset and variance are used to compute background emissions and characterize noise properties for state identification. **b**. The FRET time trace shown is produced by integrating intensities recorded at pixels in the small square neighborhood around the particle shown in the orange squares in a. Monte Carlo samples are then generated by BNP-FRET-Bin to estimate a probability distribution over states, kinetic rates, and FRET efficiencies. **c**. Distribution over state trajectories shown in pink as generated using 200 samples. Apparent FRET efficiency directly computed from data is shown in blue. **d**. Histogram over estimated state numbers. While the 3 state model is most dominant, noise intrinsic to the data does not rule out a high number of states. **e**. Apparent FRET efficiency histogram shows most of the time is spent in the intermediate FRET efficiency state. **f**. Learned FRET efficiency vs escape rate (sum of all outgoing rates from a state or inverse lifetime, λ) probability distribution. The ground truth is shown with red dots. **g**. Transition density plots showing that most transitions occur in and out of the intermediate FRET efficiency state as expected. **h-n**. Same as a-g but now for a four-state model with multiplicity of states with similar FRET efficiencies (degeneracy).

We next applied BNP-FRET-Bin to hundreds of experimental Holliday junction traces collected under tension using the MAESTRO platform [2]. In tension-free conditions, Holliday junctions populate two long-lived stacked conformations with highly heterogeneous kinetics [2] (Fig. 2a–b). Under biologically relevant, realistic four-way tension (Fig. 2c), asymmetric forces generate broad rate distributions and frequent trapping in one conformation, as evident from slow transitions at low FRET efficiency (Fig. 2e). Thermal fluctuations further alter force directions during acquisition, producing time-varying kinetic behavior; BNP-FRET-Bin captures these changes by inferring more than the two commonly-seen stacked states and resolving degeneracy (Fig. 2d). Tension also increases occupancy of the previously inaccessible intermediate open conformation (Fig. 2d), which BNP-FRET-Bin detects. As different traces visit different subsets of states, imposing a fixed number of states across traces—or estimating them trace-by-trace using histograms or BIC—would lead to systematic under- and over-fitting, as discussed above.

**Figure 2:**
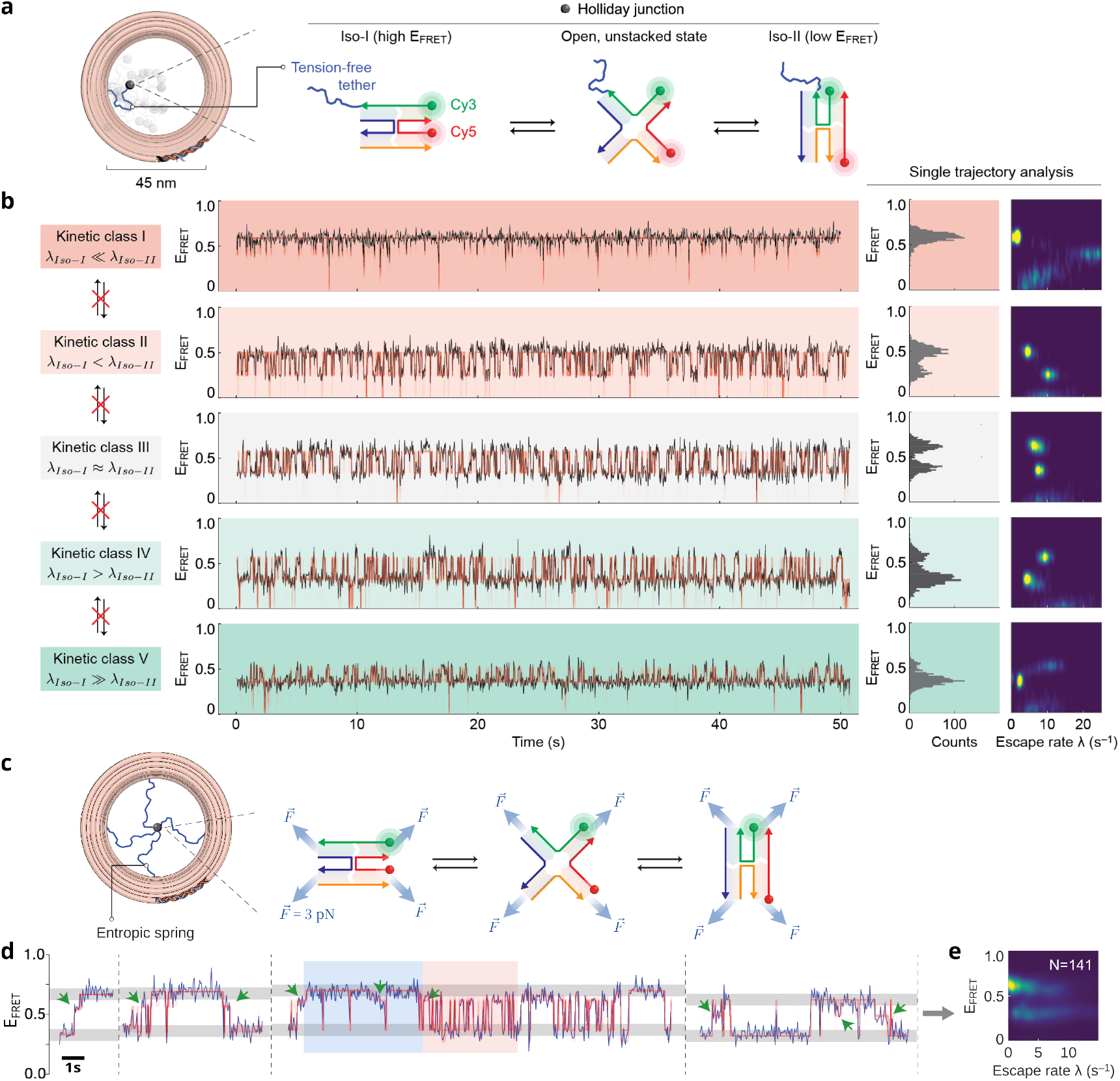
DNA Holliday junctions (HJs) under tension forces violate the “average molecule” assumption. **a**. MAESTRO platform with HJ tethered (black sphere) to a 0 pN, tension-free entropic spring (blue). HJ conformational switching between 2 states (Iso-I: E_FRET_ *∼* 0.6; Iso-II: E_FRET_ ***∼*** 0.25) monitored through a FRET pair. **b**. (Left) Heterogeneity among HJ molecules/FRET traces evident from five observed kinetic classes with distinct HJ dynamics, identified by comparing the two escape rates (inverse lifetimes) λ_*Iso−I*_ and λ_*Iso−II*_. (Middle) Representative smFRET trajectory (black) and their corresponding BNP-FRET-Bin trajectory distributions (red). (Right) E_FRET_ histograms and escape rate (λ) heatmaps obtained using BNP-FRET-Bin define the characteristic features of each kinetic class. Under unrealistic, tension-free conditions, an individual HJ molecule is not observed to switch between the five kinetic classes, indicating a rugged conformational landscape. **c**. Next, MAESTRO applies precisely controlled four-way forces on Holliday junctions via single-stranded DNA springs, simulating realistic *in vivo* environments. **d**. Sections from multiple FRET signals (blue) (separated by dashed lines) overlaid with trajectory fits (red), showing previously unseen intermediate now accessible under multiaxial forces: open unstacked conformations (green arrows). The shaded regions indicate switching between two kinetic classes with distinctly different FRET signal patterns. **e**. Resulting heatmap obtained using BNP-FRET-Bin for an ensemble of 141 HJ molecules shows a wide distribution of escape rates that would have been completely missed with existing strategies that assume an “average molecule”.

In summary, kinetic heterogeneity is expected in biomolecules due to their rugged energy landscapes. BNP-FRET-Bin offers a physically grounded, data-driven approach for such systems, particularly in the presence of degenerate FRET efficiencies, and does so without user-specified state numbers, supervised training, or ad hoc noise assumptions.

## Methods

### Lifting the “average molecule” assumption

We start by noting that smFRET tracks the motion of a biomolecule through an underlying high-dimensional, smooth conformational landscape *U* under a given data-generating process or set of experimental conditions **Λ**. For a binned smFRET experiment, these conditions may include time resolution, finite trace length limited by photobleaching, signal-to-noise, projection of all motion to one-dimensional FRET signals, detector noise, as well as environmental conditions such as temperature and viscosity. Together, these information-limiting and data-corrupting effects imply that continuous motion through the conformational landscape is not directly observed; instead, only apparent discrete jumps between long-lived conformational states are resolved. For these reasons, we restrict attention to an effective model structure ℳ that describes observable dynamics over the whole population of biomolecules in terms of an unbounded number of discrete states. Within this model, *θ* denotes the set of parameters inferred from FRET traces, such as kinetic rates and FRET efficiencies.

The fundamental uncertainty over *θ*, arising from incomplete information, data corruption, degeneracy (identical FRET signals produced by distinct conformations or experimental conditions), and discretization, is encoded in our main object of interest: the population-level distribution

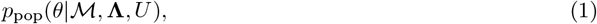

defined for a given model structure ℳ, experimental protocol **Λ**, and underlying landscape *U*.

**Note:** In the language of Bayesian inference, where a prior distribution is combined with a data likelihood to yield a posterior distribution, the expression above may naively resemble a prior, since it carries no explicit conditional dependence on a particular data realization. However, unlike the subjective modeling priors—which are analytically specified before data are observed and encode assumptions about plausible parameter values—the population distribution *p*_pop_(*θ*, |ℳ, **Λ**, *U*) above does not take a closed functional form and instead emerges after data generation. In practice, it is obtained by aggregating inference results across many independent FRET traces, such that finite data influence the distribution only through their collective ensemble.

Equivalently, the distribution above may be written as an expectation over the data-generating distribution *p*(*D*|**Λ**, *U*). Restricting attention to a fixed landscape *U*, experimental protocol **Λ**, and model structure ℳ applied to the whole population, we may write the distribution in Eq. 1 above as

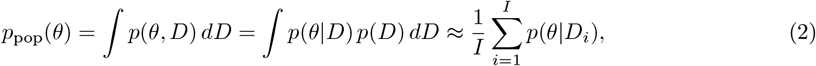

where we approximate the integral as a sum over a large number (= *I*) of independent and identically distributed (i.i.d.) FRET traces *D*_*i*_ where *i* = 1, …, *I*, drawn from the experimental data-generating distribution *p*(*D*).

Each term in the sum, *p*(*θ* | *D*_*i*_), can be computed independently by using Monte Carlo methods to generate a large number (= *J*) of samples 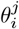, resulting in the further approximation

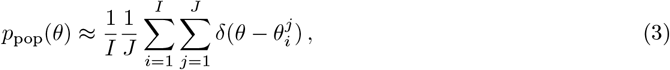

which, in principle, enables inference over a heterogeneous population of FRET traces, thereby lifting the conventional “average molecule” assumption.

A practical computational challenge nevertheless remains: although the effective model ℳ allows for an unbounded number of discrete states, any finite FRET trace can only support a finite number of parameters. A trace-by-trace Bayesian nonparametric (BNP) formulation [9], described in detail in the next subsection, provides a natural resolution. In this framework, each FRET trace activates only a finite subset of states from a conceptually infinite pool ***σ*** = {*σ*_1_, *σ*_2_, …}, with states being turned on or off as warranted by the data. Crucially, because discrete-state models are invariant under relabeling, inferred states need not correspond to unique global labels across traces: labels serve only as bookkeeping devices rather than physical identifiers. Consequently, each trace requires only a finite number of active parameter labels, and these labels may be reused across different traces without enforcing shared state identities or parameter values. Pooling Monte Carlo samples collected independently across traces then naturally aggregates over these labelings, yielding a population-level distribution over parameters without requiring explicit enumeration or alignment of the infinite state set. Operationally, this allows *θ* to be represented, for each trace, by a finite set of active parameters 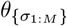, while retaining the full flexibility of the nonparametric model. The population distribution *p*_pop_(*θ*) can then be efficiently approximated as

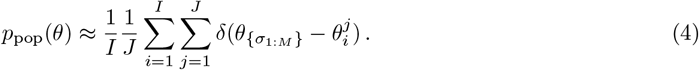

### Bayesian nonparametrics

The formulation used in our code BNP-FRET-Bin assumes a nonparametric hidden Markov model (HMM) structure to describe the time evolution of biomolecules through an infinite collection of states ***σ*** = {*σ*_1_, *σ*_2_, …} at a discrete set of equally-spaced time points **t** = {*t*_0_, *t*_1_, *t*_2_, …, *t*_*N*_} [9, 12]. *Transitions are only assumed to occur at these time points and the fixed state, which can be realized to any of the elements of* ***σ***, during a time bin (*t*_*n−*1_, *t*_*n*_) is labeled *s*_*n*_. While traditional HMM based analyses estimate transition probabilities directly, our formulation incorporates an extra step so that the transition probabilities are estimated from rates governing transition kinetics among the states via a master equation [9]. As such, BNP-FRET-Bin is designed to produce a nonparametric probability distribution, termed posterior, over an infinite collection of binary variables **b** = {*b*_1_, *b*_2_, …} that activate/deactivate states, infinite-dimensional transition rate matrix **G** governing the kinetics of the biomolecules, and an infinite collection of FRET efficiencies **E**_FRET_. Here, the nonparametric transition rate matrix **G** can be written as

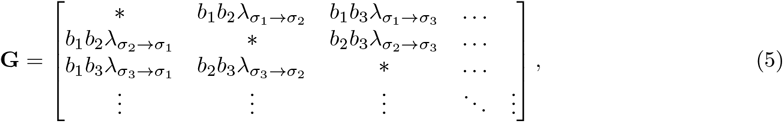

where *b*_*i*_ *∈* [0, 1] and 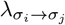 are rates for transition from state *σ*_*i*_ to *σ*_*j*_, and the *∗* in the *i*-th row of the diagonal represents the negative row-sum, 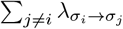, which equals the escape rate or inverse lifetime for state *σ*_*i*_. Following Bayes’ rule, we compute the posterior distribution *p*(**b, G, E**_FRET_ |***w***) as the product of the likelihood of FRET observations ***w*** = {*w*_1_, *w*_2_, …, *w*_*N*_}, *L*(***w*** | **b, G, E**_FRET_), and appropriately chosen prior probability distributions as

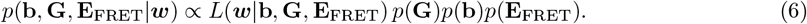

We can express the first term on the right in the proportionality above, the likelihood, for an infinite set of states [9] by observing the evolution of the state probability row-vector ***ρ*** through all the time points as

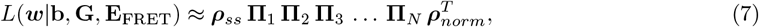

where we assume steady-state at the beginning of the FRET trace and compute the initial row-vector ***ρ***_*ss*_ by setting ***ρ***_*ss*_**G** = 0. Furthermore, **Π**_*n*_ ≡ (**Π** ⊙ **D**_*n*_) involves an element-by-element product of the transition probability matrix

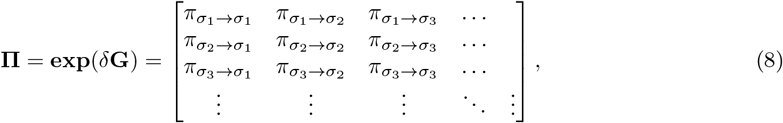

computed for a time bin of size *δ* and the measurement probability matrix whose elements are given by

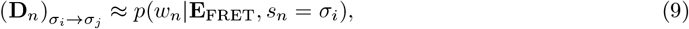

assuming the state *s*_*n*_ remains fixed during the *n*-th time bin as shown in [9].

### Likelihood

While the likelihood in Eq. 7 above integrates over all possible state trajectories, computational cost associated with a large number of matrix exponentials can be very high. Therefore, instead of repeatedly computing the whole likelihood to sample each parameter of interest, we integrate by sampling the state trajectory ***s*** = {*s*_1_, *s*_2_, …, *s*_*N*_} itself. That is, we rewrite the likelihood as

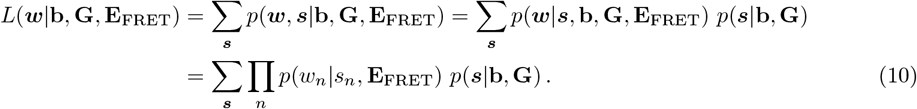

Now, computing these measurement probabilities above in Eq. 9 in order to obtain the likelihood is challenging because of the many unobserved intermediate random variables that may assume values from a wide range of possibilities to result in exactly the same set of observations. For example, when using sCMOS cameras to detect signal, Gaussian noise is added to the signal at readout [13] which adds uncertainty to the number of donor/acceptor photons entering the detectors. Similarly, the photon shot noise adds uncertainty to the underlying FRET efficiency associated with a state. Furthermore, in the case of EMCCD cameras, an intermediate electron multiplication stage adds Gamma distributed noise [14] and uncertainty to the detected number of photons. More formally, we must propagate all this uncertainty into the measurement probabilities *p*(*w*_*n*_ | *s*_*n*_, **E**_FRET_) by integrating over all possible values of the unobserved random variables including the number of excitation laser photons absorbed by the donor 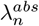 in each time bin related to the average excitation rate λ^*ex*^, donor and acceptor channel photon emissions per time bin, 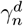 and 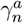, respectively, and multiplied electrons in the donor and acceptor channels per time bin, 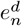 and 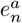, respectively, if an EMCCD camera is used. That is, for sCMOS cameras,

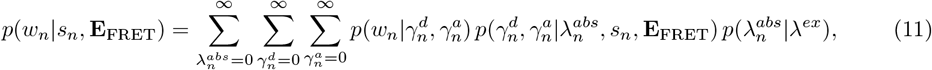

and for EMCCD cameras,

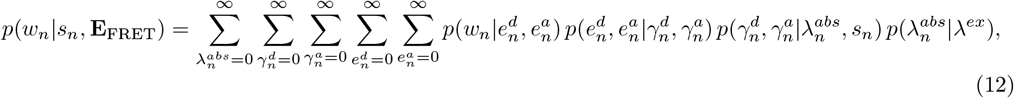

where the explicit expressions for probabilities are

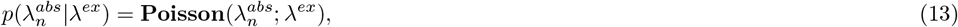

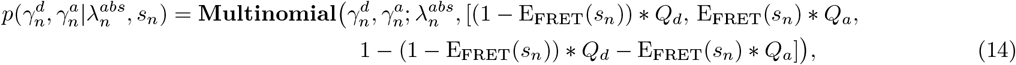

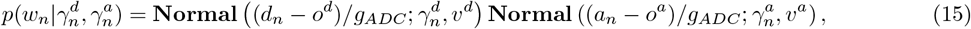

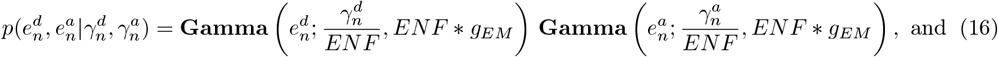

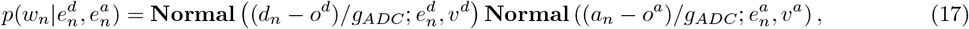

where square brackets represent probability vectors, *o*^*d*^ and *o*^*a*^ are bias offsets for the donor and acceptor channels, respectively, *g*_*ADC*_ and *g*_*EM*_ are analog-to-digital conversion factor and electron multiplier gain, respectively, and *Q*_*a*_ and *Q*_*d*_ are quantum yields of donor and acceptor, respectively. Furthermore, due to the stochastic nature of electron multiplication itself in EMCCD cameras, the excess noise factor, *ENF* = 2.0 *−* 1*/g*_*EM*_, affects the electron multiplication significantly when gain is high and readout noise becomes relatively negligible.

These integrations in Eqs. 10-12 are not easily amenable to analytical computations and therefore we integrate by Monte Carlo as we describe next.

### Sampling

Although the complex hierarchy above with a large number of nested sums and products in the probability expressions appear intimidating, a nice property of probabilities is that all conditional dependencies, correlations, and all uncertainty from every source are encoded in a single full joint distribution over all unknown variables [15, 12],

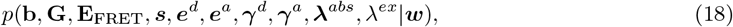

where the boldface variables ***e***^*d*^, ***e***^*a*^, ***γ***^*d*^, ***γ***^*a*^, and **λ**^*abs*^ collect the previously discussed latent variables at each time point *t*_*n*_ succinctly. This simple property implies that sampling from the joint posterior distribution automatically propagates the uncertainty into the individual probability distribution over any variable once we marginalize (integrate) over all others. Most importantly, the joint posterior above is amenable to Gibbs sampling, where we generate samples for a variable sequentially and iteratively, assuming all other variables are constant [15, 12], as we now show.

After appropriately initializing all variables, the following Gibbs sampling strategy is employed in BNP-FRET-Bin to sequentially and iteratively generate samples:

1. Truncate the infinite-dimensional model for computational reasons only to finite but large number of states *M*. Then jointly sample loads and state trajectory to avoid inconsistency between the active states and states being explored in the trajectory. To accomplish this, two trajectories corresponding to the two values of the binary variable *b*_*i*_ are proposed 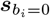 and 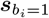using the forward filtering backward sampling algorithm [12] as

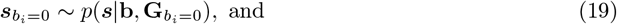

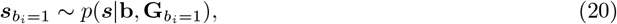

where we have ignored variables that do not directly inform the trajectories. Likelihoods are then computed for both trajectories and then conditional posteriors are computed using the Bernoulli prior for *b*_*i*_ as

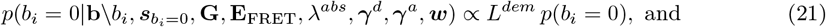

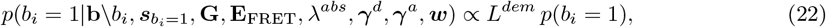

where the prior with expected number of states *M*_*exp*_ is given as

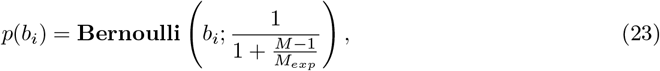

and suppresses activation of new states with the probability shown after the semi-colon to avoid over-fitting. For typical FRET traces, we find *M* = 20 and *M*_*exp*_ = 2 works well. The new value of *b*_*i*_ can now be sampled from a Categorical distribution as

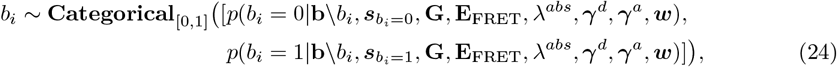

and the trajectory associated with the sampled value of *b*_*i*_ is accepted as the new trajectory. In the last equation above, element appearing immediately to the right of “\” is excluded from the set on the immediate left.
2. Sample kinetic rates

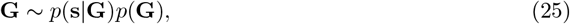

Where

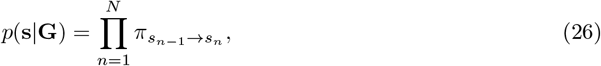

since only the kinetic rates affect the state trajectory. Furthermore, the prior *p*(**G**) can be chosen as a product of Gamma distributions over each rate to enforce positivity.
3. Sample FRET efficiencies

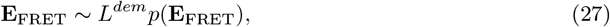

where Beta distributions can be used as priors to enforce the finiteness of the domain, 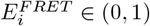
4. Sample donor excitation rate

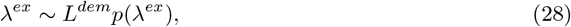

where Gamma distribution can be used for the prior to enforce positivity.
5. Sample donor photon absorptions

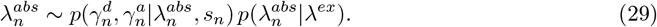
6. Sample donor and acceptor emissions

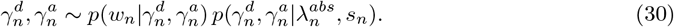
7. Sample multiplied electrons in donor and acceptor channels (in the case of EMCCD cameras)

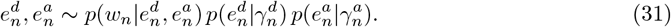

In cases where the conditional posteriors are not amenable to direct sampling, we employ techniques such as Metropolis-Hastings to generate samples. Once enough uncorrelated MCMC samples have been collected, they can be used to compute statistical quantities such as mean, uncertainties, and confidence intervals if desired.

## Acknowledgments

RFH acknowledges support from NSF (MCB 2341002). SP acknowledges support from NIH (R01GM134426, R01GM130745, R35GM148237), and US Army (ARO W911NF-23-1-0304). GBMW is supported by an AHA predoctoral fellowship (23PRE1029870). We acknowledge the use of facilities within the Eyring Materials Center as well as Phoenix and Sol clusters at Arizona State University.

## Bibliography

[1] D. Nettels, N. Galvanetto, M. T. Ivanović, M. Nüesch, T. Yang, and B. Schuler. “Single-molecule FRET for probing nanoscale biomolecular dynamics”. Nature Reviews Physics 6 (2024), p. 587.

[2] G. B. M. Wisna, A. Saurabh, D. Karna, R. Sasmal, P. Chopade, S. Pressé, and R. F. Hariadi. “Multi-axial DNA origami force spectroscopy reveals hidden dynamics of Holliday junctions”. bioRxiv (2025).

[3] E.M. López-Alfonzo, A. Saurabh, S. Zarafshan, S. Pressé, and A. Martin. “Substrate-interacting pore loops of two ATPase subunits determine the degradation efficiency of the 26S proteasome”. bioRxiv (2023).

[4] M. Götz and et al. “A blind benchmark of analysis tools to infer kinetic rate constants from single-molecule FRET trajectories”. Nature Communications 13 (2022), p. 5402.

[5] S. Wanninger, P. Asadiatouei, J. Bohlen, C.-B. Salem, P. Tinnefeld, E. Ploetz, and D. C. Lamb. “Deep-LASI: deep-learning assisted, single-molecule imaging analysis of multi-color DNA origami structures”. Nature Communications 14 (2023), p. 6564.

[6] J. Thomsen, M. B. Sletfjerding, S. B. Jensen, S. Stella, B. Paul, M. G. Malle, G. Montoya, T. C. Petersen, and N. S. Hatzakis. “DeepFRET, a software for rapid and automated single-molecule FRET data classification using deep learning”. Elife 9 (2020), e60404.

[7] A. Saurabh, L. W. Q. Xu, and S. Pressé. “On the statistical foundation of a recent single molecule FRET benchmark”. Nature Communications 15 (2024), p. 3627.

[8] A. Bandyopadhyay and M. P. Goldschen-Ohm. “Unsupervised selection of optimal single-molecule time series idealization criterion”. Biophysical Journal 120 (2021), p. 4472.

[9] A. Saurabh, M. Fazel, M. Safar, I. Sgouralis, and S. Pressé. “Single-photon smFRET. I: Theory and conceptual basis”. Biophysical Reports 3 (2023).

[10] I. Sgouralis, S. Madaan, F. Djutanta, R. Kha, R. F. Hariadi, and S. Pressé. “A Bayesian Nonparametric Approach to Single Molecule Förster Resonance Energy Transfer”. The Journal of Physical Chemistry B (2019), p. 675.

[11] A. Saurabh. https://github.com/LabPresse/BNP-FRET-Binned.

[12] S. Pressé and I. Sgouralis. Data modeling for the sciences: applications, basics, computations. Cambridge University Press, 2023.

[13] B. Mandracchia, X. Hua, C. Guo, J. Son, T. Urner, and S. Jia. “Fast and accurate sCMOS noise correction for fluorescence microscopy”. Nature Communications 11 (2020), p. 1.

[14] M. Hirsch, R. J. Wareham, M. L. Martin-Fernandez, M. P. Hobson, and D. J. Rolfe. “A stochastic model for electron multiplication charge-coupled devices–from theory to practice”. PLOS ONE 8 (2013), p. 1.

[15] C. M. Bishop and N. M. Nasrabadi. Pattern recognition and machine learning. Vol. 4. Springer, 2006.

